# Genomics of expanded avian sex chromosomes shows that certain chromosomes are predisposed towards sex-linkage in vertebrates

**DOI:** 10.1101/585059

**Authors:** Hanna Sigeman, Suvi Ponnikas, Pallavi Chauhan, Elisa Dierickx, M. de L. Brooke, Bengt Hansson

## Abstract

Sex chromosomes have evolved from the same autosomes multiple times across vertebrates, suggesting that certain genomic regions are predisposed towards sex-linkage. However, to test this hypothesis detailed studies of independently originated sex-linked regions and their gene content are needed. Here we address this problem through comparative genomics of birds where multiple chromosomes appear to have formed neo-sex chromosomes: larks (Alaudidae; Sylvioidea). We detected the largest known avian sex chromosome (195.3 Mbp) and show that it originates from fusions between (parts of) four avian chromosomes (Z, 3, 4A and 5). We found evidence of five evolutionary strata where recombination has been suppressed at different time points, and that these time points correlate with the level of Z–W gametolog differentiation. We show that there is extensive homology to sex chromosomes in other vertebrate lineages: three of the fused chromosomes (Z, 4A, 5) have independently evolved into sex chromosomes in fish (Z), turtles (Z, 5), lizards (Z, 4A) and mammals (Z, 4A). Moreover, we found that the fourth chromosome, chromosome 3, was significantly enriched for genes with predicted sex-specific functions. These results support a key role of chromosome content in the evolution of sex chromosomes in vertebrates.

## 1. INTRODUCTION

Sex chromosomes have evolved from autosomes many times independently across the tree of life. The most generic hypothesis for the evolution of sex chromosomes invokes a selective advantage of linkage between sex-determining and sexually antagonistic genes (Haldane 1922; Fisher 1931; Lewis & John 1968; D. Charlesworth & B. Charlesworth 1980). Accordingly, this hypothesis suggests that the chromosomes harbouring these genes should frequently be involved in the formation, transition and turnover of sex chromosomes (Bachtrog et al. 2011; Ross et al. 2009; O’Meally et al. 2012). Indeed, there are a limited number of genes across vertebrates that have taken on the sex-determining role, and evidence is accumulating that the chromosomes on which these reside have evolved into sex chromosomes several times independently in different lineages (reviewed in O’Meally et al. 2012; Ezaz et al. 2006; Ezaz et al. 2016). Furthermore, it has been suggested that some autosomes fuse with existing sex chromosomes into neo-sex chromosomes more easily than others due to their gene content (Kitano et al. 2009; Ross et al. 2009; Pala, Hasselquist, et al. 2012; Kitano & Peichel 2011; Yoshida et al. 2014; O’Meally et al. 2012). The importance of chromosome content for the evolutionary dynamics of sex chromosomes is, however, still poorly supported due to a limited number of well-studied origins of sex-linkage.

The sex chromosomes of birds are highly stable with a Z chromosome size of approximately 73 Mbp (Ellegren 2010). One of the few known exceptions to this stability is the neo-sex chromosome of passerines in the superfamily Sylvioidea, which has been formed by a fusion between half of chromosome 4A (according to the zebra finch *Taeniopygia guttata* nomenclature) and the ancestral sex chromosome (Pala, Naurin, et al. 2012; Pala, Hasselquist, et al. 2012; Sigeman et al. 2018). Moreover, two independent findings suggest that at least some species of larks (Alaudidae), a family within Sylvioidea, have acquired autosome– sex chromosome fusions in addition to the one including parts of 4A. Firstly, heavily enlarged sex chromosome karyotypes were found in the bimaculated lark (*Melanocorypha bimaculata*) and the horned lark (*Eremophila alpestris*; Bulatova 1981), and, secondly, genetic markers located on chromosome 3 and 5 were found to have sex-specific inheritance in the Raso lark (*Alauda razae*; Brooke et al. 2010).

Here, we use comparative genomics to study this neo-sex chromosome system across three Alaudidae species – two *Alauda* species (Raso lark and Eurasian skylark *A. arvensis*) and one *Eremophila* species (horned lark) – and their sister species in the family Panuridae (the bearded reedling *Panurus biarmicus*). We use whole-genome sequence data of these species to characterise in detail which genomic regions have become fused to the sex chromosomes, and use phylogenetic information to determine the appearance and age of several evolutionary strata where recombination has been suppressed at different points in time (*sensu* Lahn & Page 1999). Then, we test whether the age of the strata explains the degree of Z-W divergence of gametologous genes, as predicted by sex chromosome theory (Rice 1994; Lahn & Page 1999). Next, we test the hypothesised importance of chromosome content for the formation of sex chromosomes by analysing whether the fused chromosomal regions are enriched for genes with sex-specific functions. Lastly, we evaluate signs of repeated sex chromosome evolution by searching for homologies between the fused sex chromosomes in larks and the sex chromosomes of other vertebrate lineages. Our study supports a key role of chromosome content in the evolution of sex chromosomes and highlights the importance of studying independently originated sex chromosomes for understanding how and why vertebrate sex chromosomes evolve.

## 2. MATERIALS & METHODS

### 2.1 Sequence data

We extracted DNA from blood samples using a phenol–chloroform protocol (Sambrock & W Russel, 2001) of one female and one male individual from each of our four study species: Raso lark (*Alauda razae*, from Cape Verde), Eurasian skylark (*A. arvensis cantarella*, from Italy), horned lark (*Eremophila alpestris flava*, from Sweden) and bearded reedling (*Panurus b. biarmicus*, from Sweden). According to *cytochrome b* sequence data, the two *Alauda* species diverged c. 6 million years ago (Mya), *Alauda* and *Eremophila* c. 14 Mya (Alström et al. 2013), and Alaudidae and Panuridae c. 17 Mya (Moyle et al. 2016). All samples were collected non-destructively and with permission from the relevant authorities (Direcção Geral do Ambiente, Cape Verde, and Malmö/Lund Ethical Committee for scientific work on animals, Sweden, no. 17277-18). DNA was sequenced with Illumina HiSeqX (150 bp, paired-end) by SciLifeLab Sweden.

### 2.2 *De novo* assembly and mapping

To obtain reference genomes, we created *de novo* genome assemblies from the male sequence data for each of the study species. The resequencing data was trimmed using nesoni clip (https://github.com/Victorian-Bioinformatics-Consortium/nesoni) with a minimum read quality of 20 and minimum read length of 20 bp. The trimmed reads were then assembled with Spades v3.5.0 (Bankevich et al. 2012) using 5 different kmer lengths (21, 33, 55, 77 and 127) and the setting “careful”. Scaffolds shorter than 1 kbp were discarded. Quality statistics from the assemblies were calculated using Quast v4.5.4 (Gurevich et al. 2013). Assemblies with N50 over 50 kbp were kept for further analysis. This threshold meant keeping the assemblies of the Raso lark (N50=103 kbp, 28304 scaffolds) and the bearded reedling (N50=68 kbp, 36455 scaffolds), while the assemblies for the Eurasian skylark (N50=8 kbp, 256822 scaffolds) and horned lark (N50=22 kbp, 109088 scaffolds) were discarded.

Reads from all of the samples (n = 8) were cleaned for adaptor sequences with Trimmomatic v.0.3.6 (Bolger et al. 2014) using the adaptor file TruSeq3-PE and options seedMismatches: 2, palindromeClipThreshold: 30 and simpleClipThreshold: 10. Trimming of low-quality bases was done using a quality threshold of 15 from the leading end and 30 from the trailing end. The reads were further trimmed for a minimum quality of 20 over sliding windows of 4 bp. Lastly, any reads shorter than 90 bp were excluded from further analyses. The number of remaining reads in our samples ranged from 176 to 404 million. The samples were quality checked using fastqc v0.11.7 (https://www.bioinformatics.babraham.ac.uk/projects/fastqc/).

The male and female samples of Raso lark and bearded reedling were aligned to their respective genome assemblies, while the Eurasian skylark and horned lark individuals were aligned to the genome assembly of their closest relative, the Raso lark. The alignment was done with bwa mem v0.7.17 (Li & Durbin 2009), marking shorter split hits as secondary (option -M) for downstream compatibility. The aligned reads were sorted with samtools v1.7 (Li et al. 2009), and duplicated reads were removed using picardtools v2.18.0 (http://broadinstitute.github.io/picard). Assembly and alignment statistics are provided in Suppl. Tables S1,2.

### 2.3 Chromosome anchoring

The scaffolds in the genome assemblies were grouped into different chromosomes and ordered into chromosome-level using the genome assembly of the zebra finch (*Taeniopygia guttata*; (Warren et al. 2010). The zebra finch genome assembly taeGut.3.2.4 was downloaded from Ensembl (Cunningham et al. 2015) and transformed into a database using the last v876 (Kielbasa et al. 2011) program lastdb. The two genome assemblies, of the Raso lark and bearded reedling, were aligned to the zebra finch genome using the program lastal and converted to psl format using the script maf-convert, both from the same software suite last v876. From there, we extracted chromosome anchoring coordinates based on the longest match to the zebra finch genome for each 5 kbp window in the Raso lark and bearded reedling assemblies, with a minimum requirement of 500 matching base pairs per 5 kbp window. The assembly positions from the output files of the coverage and single nucleotide variant (SNV) analyses were then translated to the starting positions of the match to the zebra finch genome assembly (see section below).

### 2.4 Identification of sex-linked genomic regions

We identified sex-linked regions using two different kinds of genomic signatures: (i) differential mapping success in males and females (i.e. sex-specific genome coverage), and (ii) an excess or deficit of female-specific genetic variation. Sex chromosomes almost invariably evolve recombination suppression in the heterogametic sex so that regions that have been sex-linked for a long time (such as the sex chromosomes that formed in the ancestor of all birds) will show pronounced sequence divergence and degeneration in the non-recombining chromosome (the W in birds; Zhou et al. 2014). The regions belonging to the ancestral sex chromosomes can thus be identified by lower female coverage, and fewer female-specific genetic variants compared to males. This is because reads from the female-specific W-chromosome will either not map to the male reference genome due to substantial differentiation between the Z and W or because of deletions on the W chromosome. More-recently formed sex-linked regions may be identified by lower mapping success in females, although we expect a subtler difference as the W-linked genomic region may not have yet developed substantial differentiation from the Z-linked homologous region. Here, we also expect a higher amount of female-specific mutations compared to males, due to differentiation between the Z and W sex chromosome copies.

To uncover differential mapping success in males and females for each species, we parsed the alignment files for reads with more than two mismatches and calculated genome-wide coverage for 5 kbp windows with bedtools v2.71.1 (Quinlan & Hall 2010). All genome coverage values were normalised between the female and the male sample, based on the number of reads in the trimmed and adaptor-free fastq files. To identify sex-linked regions, we binned the female-to-male coverage values for every 1 Mbp genomic region and extracted the mean value from each bin. Following discovery of sex-linked genomic regions, we performed another normalisation step by dividing the median female-to-male coverage ratio for each 5 kbp window by the genome-wide median female-to-male coverage ratio counting only chromosomes without sex-linked regions. We then binned the data into 0.1 Mbp windows and calculated the mean female-to-male coverage ratio within each bin.

To analyse female-specific variation, we called variants in the alignment files (with all mismatches allowed) with freebayes v1.1.0 for each species separately (n = 2 in each analysis) using freebayes-parallel (Garrison & Marth 2012) and parallel v20180322 (Tange 2018). The output was then parsed for any SNP that had been marked with a flag other than PASS (--remove-filtered-all), a minimum quality of 20 and minimum depth of 3x using vcftools v0.1.15 (Danecek et al. 2011). Private alleles (minor alleles occurring only in one sample in a heterozygous state) were extracted with vcftools using option --singletons. We calculated the difference between the number of female-specific private alleles and male-specific private alleles for each 5 kbp window and extracted the average difference across 1 Mbp and 0.1 Mbp windows.

### 2.5 Analysis of divergence between gametologous genes

We used the whole-genome synteny aligner program SatsumaSynteny v. 2.0 (Grabherr et al. 2010) to align the Raso lark assembly to the zebra finch assembly (taeGut.3.2.4), and then used kraken (Zamani et al. 2014, downloaded 18 June 2018) to make a lift-over of the zebra finch annotations to the Raso lark assembly. Of the 18204 transcripts and 17488 genes in the zebra finch annotation, 14466 transcripts (79 %) from 13764 genes (79 %) were annotated in the Raso lark. We used Freebayes v.1.1.0 (Garrison & Marth 2012) (--report-monomorphic) to call variants for every base pair within all exons in the Raso lark based on the genome coordinates from the lift-over.

We *in silico* extracted gametologous gene sequences from sex-linked regions using an in-house script (code provided as Supplementary Code S1; general methodology described in (Sigeman et al. 2018) based on the genotypes of the female and male samples. The script uses sex-specific genotype information to phase the data into a Z and W sequence. We replaced a site by “N” if it had a quality score or sequence depth below 20 or if the genotype in either sample was not called. Any site without variants in either sample was extracted as such, and remaining variants between the male and female were phased based on sex-specific allele compositions provided in Suppl. Table S3.

To confirm that both Z and W gametologs were present in the data, we calculated genome coverage values for every exon using bedtools v.2.27.1 multicov (Quinlan & Hall 2010) and normalised the values between the two samples of each species. Exons with female coverage less than 75 % of the male sample were masked with N:s, as this suggests absence of a W gametolog.

We used TransDecoder v3.0.1 (https://github.com/TransDecoder/TransDecoder/) to find the longest open reading frames for each of the sequences (--retain_pfam_hits, --retain_blastp_hits, --single-best-orf). Orthologous genes from the zebra finch, collared flycatcher (*Ficedula albicollis*) and chicken (*Gallus gallus*) were downloaded through BioMart (database: Ensembl Genes 93), and the longest transcript from the flycatcher and chicken corresponding to the zebra finch transcripts in the annotation was added as additional sequences. These gene sequences were not used in further analyses but aided in evaluating the quality of the alignments. The sequences were codon-aware aligned using *Prank* v.150803 (Loytynoja 2014). Each N in the sequences was transformed into “-”, and all sites including this character in any sequence were then removed using gblocks v0.91b (Castresana 2000). We calculated pairwise substitution rates between Z–W gametologs using codeml (estimated kappa and codon frequency F3X4) from the PAML v4.9 package (Yang 2007). Gene sequences longer than 500 bp, and where the Z and W sequences within a species had a synonymous substitution rate (dS) value above 0.01, were kept for further analysis. We used Kruskal-Wallis tests to investigate potential differences in sequence divergence between the identified sex-linked genomic regions (or strata, see below) as the data were not normally distributed. All values were log2-transformed prior to these tests to fulfil the criteria of similar distributions between groups and a low number (0.00001) was added to all synonymous substitution rate (dN) values as some contained zeros. Significance levels of differences between groups were calculated using pairwise Wilcoxon tests with Benjamini & Hochberg adjusted p values. Furthermore, we used ordered heterogeneity tests (OH; Rice & Gaines 1994) to evaluate if the difference between Z–W nucleotide divergences between groups correlated with relative timing of sex-linkage between the evolutionary strata. The OH value r_s_p_c_ was calculated by multiplying the complement of the p value from the Kruskal-Wallis tests (1 – p value) with the Spearman Rank rho value from the median nucleotide divergence estimates for each stratum correlated with the order in which they appear based on the phylogeny of the studied species. The relative timing of sex-linkage of stratum 5a and 5b (see below) could not be distinguished using our data and were thus given the same rank. P values were extracted from the supplied figures in Rice & Gaines (1994), and multiplied by two for two-tailed hypothesis testing.

### 2.6 Gene enrichment analysis

We downloaded Gene Ontogeny (GO) annotations for all the genes in the zebra finch genome from Ensembl BioMart (taeGut3.2.4; accessed on 12 September 2018) and parsed the file for GO term names with the following pattern matches: “sperm”, “ovarian”, “gonad”, “estrogen”, “testosterone”, “sex differentiation”, “sex determination”, “sexual characteristics”, “sexual reproduction” and “oogenesis”. The zebra finch gene annotation had in total 323 genes matching to any of these patterns. We then counted the number of genes matching any of these terms for i) the sex-linked region for each chromosome separately, and ii) within each of the identified evolutionary strata (see below). We performed two-tailed binomial tests to see if these genomic regions contained either more or fewer sex-related genes than would be expected if the genes were randomly distributed across the genome. We calculated the proportion of the genome that made up each studied sex-linked region (1.253 Gbp is the full genome size of the zebra finch genome assembly used) and tested if the region contained a higher or lower proportion of sex-related genes than would be expected based on this probability. The p values for the two categories of tests (i, ii) were adjusted using the Benjamini & Hochberg method.

## 3. RESULTS

### 3.1 Identification of sex-linked genomic regions

A genome-wide scan for sex-specific genetic variation (measured as the average difference in number of female and male private SNVs per Mbp), and sex-specific genome coverage (measured as the average female-to-male read coverage ratio per Mbp), revealed that four chromosomes stood out from the autosomal pattern across the species (Figure 1; Suppl. Table S4). First, the ancestral Z chromosome showed substantially lower female coverage, and no or a moderately lower amount of female-specific genetic variation, in all four species (Figure 1). Secondly, the first half of chromosome 4A showed substantially higher number of female-specific SNVs and moderately lower female coverage in all species. Thirdly, a substantially higher amount of female-specific SNVs and no or moderate sex-specific coverage were found over varying extents of chromosome 3 and 5 in the four species – small parts of chromosome 3 in the bearded reedling, slightly larger parts of chromosome 3 in the horned lark, and the greater part of chromosome 3 and 5 in the Raso lark and the Eurasian skylark (Figure 1). These results support the existence of multiple autosome–sex chromosome fusions and several evolutionary strata where recombination has been suppressed at different points in time across the avian and the lark phylogeny.

**Figure 1.**
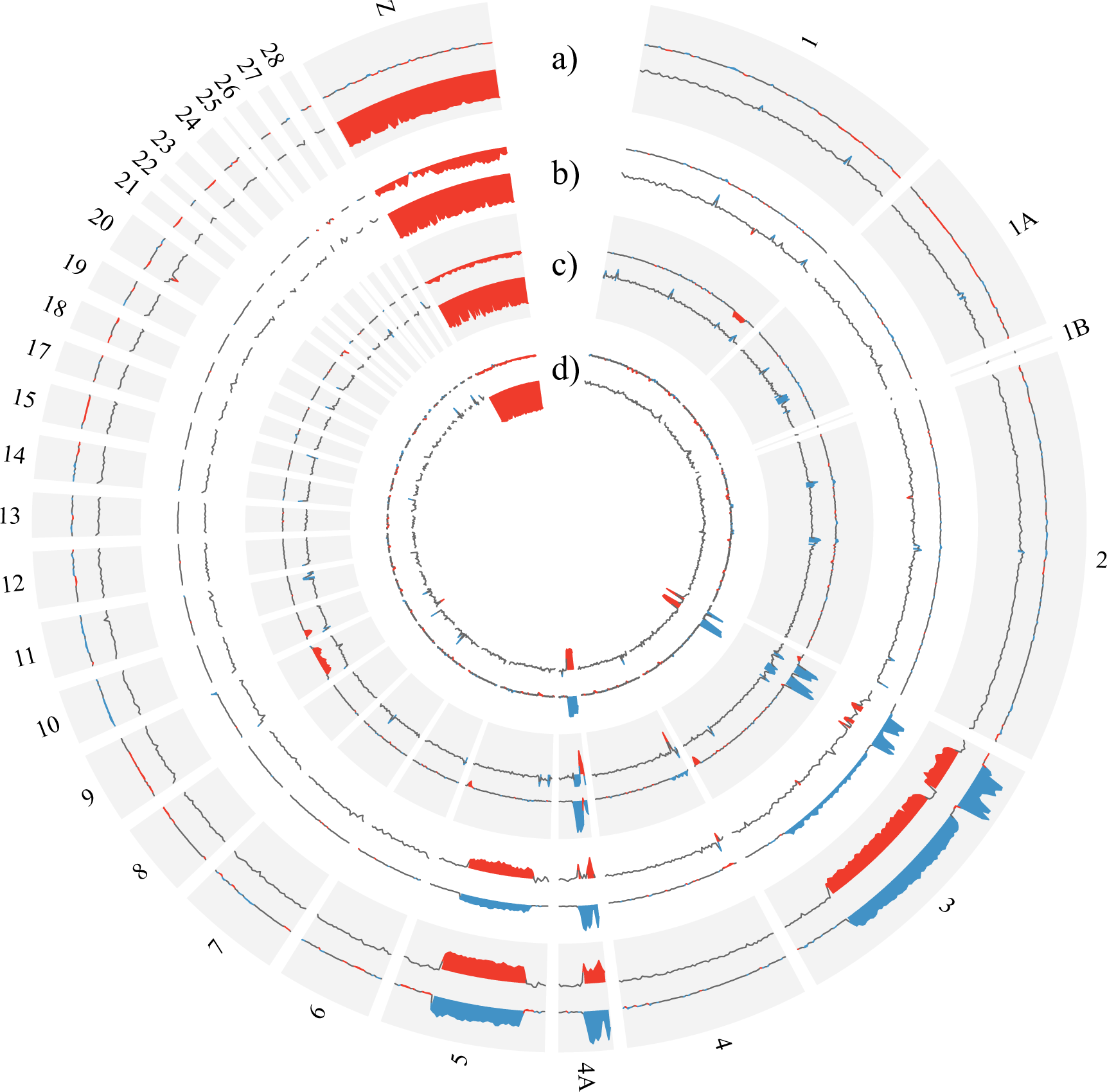
Genome-wide distribution of female-to-male difference in number of private single nucleotide variants (SNVs), and female-to-male coverage ratio, in the four study species. The background colours grey and white separate the data for each of the four species which are shown in the following order starting from the outer ring: a) Raso lark, b) Eurasian skylark, c) horned lark and d) bearded reedling. Within each ring, the outer line shows the average difference in number of private SNVs between females and males across 1 Mbp windows (with values > 500 in blue and values < -500 in red), and the inner line shows the average female-to-male coverage ratio across 1 Mbp windows (with values > 1.1 in blue and values < 0.9 in red). Chromosome-wide averages are provided in Suppl. Table S4.

To define the evolutionary strata more precisely, and to estimate their age, we assessed (i) the genomic signatures of sex-linkage (i.e., sex-specific genetic variation and genome coverage as above) at the scale of 0.1 Mbp regions in each species (Table 1; Suppl. Figure S1), and (ii) the most parsimonious order of emergence based on dated phylogenies (Figure 2; Alström et al. 2013, Cortez et al. 2014, Moyle et al. 2016). Stratum 1 corresponds to the entire ancestral Z chromosome (72.9 Mbp), which was formed c. 140 Myr ago (Cortez et al. 2014), and stratum 2 corresponds to the previously known Sylvioidea neo-sex chromosome, i.e. the first 9.6 Mbp of chromosome 4A (Pala et al. 2012), which was formed c. 21-19 Myr ago when Sylvioidea split from other passerines (Moyle et al. 2016). For both these strata, all four study species showed clear signatures of sex-linkage (Table 1; Suppl. Figure S1). Likewise, all four study species showed sex-specific genetic variation and coverage in the regions between 8.4–10.4 and 18.1–24.1 Mbp on chromosome 3 (with a total size of 8.0 Mbp): this constitutes stratum 3, which is the earliest stratum being unique to the Alaudidae/Panuridae clade. The age of stratum 3 is defined by the split of Alaudidae/Panuridae from other Sylvioidea families c. 19-17 Myr ago (Moyle et al. 2016). The next stratum, stratum 4, has a size of 3.6 Mbp and spans the region between 10.4–14.0 Mbp on chromosome 3. It occurs in all three lark species, but not in the bearded reedling, which gives an estimated age of c. 17-14 Myr as defined by the split between Alaudidae and Panuridae (Alström et al. 2013). Finally, the Raso lark and the Eurasian skylark had two additional strata on chromosome 3 (stratum 5a: 64.9 Mbp spanning the regions between 5.8–8.4, 14.0-18.1 and 29.8-88.0 Mbp) and chromosome 5 (stratum 5b: 36.3 Mbp, the region between 9.1-45.4 Mbp), respectively, that were not shared with the other two species. These strata form the youngest sex chromosome layer, estimated to be c. 14-6 Myr based on the split between the horned lark and the *Alauda* larks (Alström et al. 2013). Together, these sex-specific regions on chromosome Z, 4A, 3 and 5 amounts to 195.3 Mbp of the genome of the Raso lark and Eurasian skylark (16.3% of the genome based on a genome size estimation of 1.2 Gbp) (Table 1; Suppl. Figure S1). The horned lark had 94.1 Mbp (7.8%), and the bearded reedling 90.5 Mbp (7.5%), sex-linked genomic material (Table 1; Suppl. Figure S1). Figure 2a shows a summary of the chromosomal position of the evolutionary strata and their most parsimonious order of appearance based on presence and absence in the studied species (also shown are silhouettes indicating vertebrate groups where homologs to these chromosomes act as sex chromosomes; see Discussion). Figure 2b summarises the timings of sex-linkage based on dated phylogenies (Cortez et al. 2014, Alström et al. 2013, Moyle et al. 2016).

**Table 1.** Sex-linked regions in each of the four species. The strata are numbered according to the most parsimonious order of emergence based on phylogenetic analyses. Stratum 1 (homologous to chromosome Z in zebra finch) acts as the sex chromosome in all birds, and sex-linkage of stratum 2 (chromosome 4A in zebra finch) is common for all birds belonging to the superfamily Sylvioidea, which includes larks and the bearded reedling. Stratum 3 was seen to be sex-linked in all four studied species and is therefore older than stratum 4, which appears as sex-linked in all larks but not the bearded reedling. Stratum 5a and 5b appear as sex-linked in the Raso lark and Eurasian skylark, but not in horned lark or bearded reedling, and is therefore the youngest stratum but here divided into two substrata as they originated from different autosomes. The table provides mean values for the male-to-female coverage ratio (coverage) and female-to-male difference in number of private SNVs across 0.1 Mbp windows for each stratum. Bearded reedling and horned lark have no sex-linkage for stratum 5a and 5b, and bearded reedling not for stratum 4, and are therefore marked with NA in addition to the corresponding values.

| Stratum | Chromosome | Genomic region (Mbp) | Stratum Size (Mbp) |  | Raso lark | Eurasian skylark | Horned lark | Bearded reedling |
| --- | --- | --- | --- | --- | --- | --- | --- | --- |
| 1 | Z | 0-72.9 | 72.9 | Coverage | 0.52 | 0.55 | 0.54 | 0.52 |
|  |  |  |  | SNV | -17.77 | -992.10 | -375.85 | -160.48 |
| 2 | 4A | 0-9.6 | 9.6 | Coverage | 0.73 | 0.89 | 1.06 | 0.66 |
|  |  |  |  | SNV | 2957.35 | 2227.36 | 2518.46 | 2413.16 |
| 3 | 3 | 8.4-10.4,<br>18.1-24.1 | 8.0 | Coverage | 0.74 | 0.95 | 1.20 | 0.68 |
|  |  |  |  | SNV | 3386.68 | 2435.65 | 3404.79 | 2787.17 |
| 4 | 3 | 10.4-14.0 | 3.6 | Coverage | 0.69 | 0.84 | 0.97 | NA (1.00) |
|  |  |  |  | SNV | 3019.06 | 2161.91 | 2512.71 | NA (-10.54) |
| 5a | 3 | 5.8-8.4,<br>14.0-18.1,<br>29.8-88.0 | 64.9 | Coverage | 0.74 | 0.98 | NA (1.00) | NA (0.99) |
|  |  |  |  | SNV | 1785.39 | 451.48 | NA (1.73) | NA (1.77) |
| 5b | 5 | 9.1-45.4 | 36.3 | Coverage | 0.69 | 0.85 | NA (1.00) | NA (0.99) |
|  |  |  |  | SNV | 1952.04 | 879.14 | NA (20.19) | NA (-37.92) |

**Figure 2.**
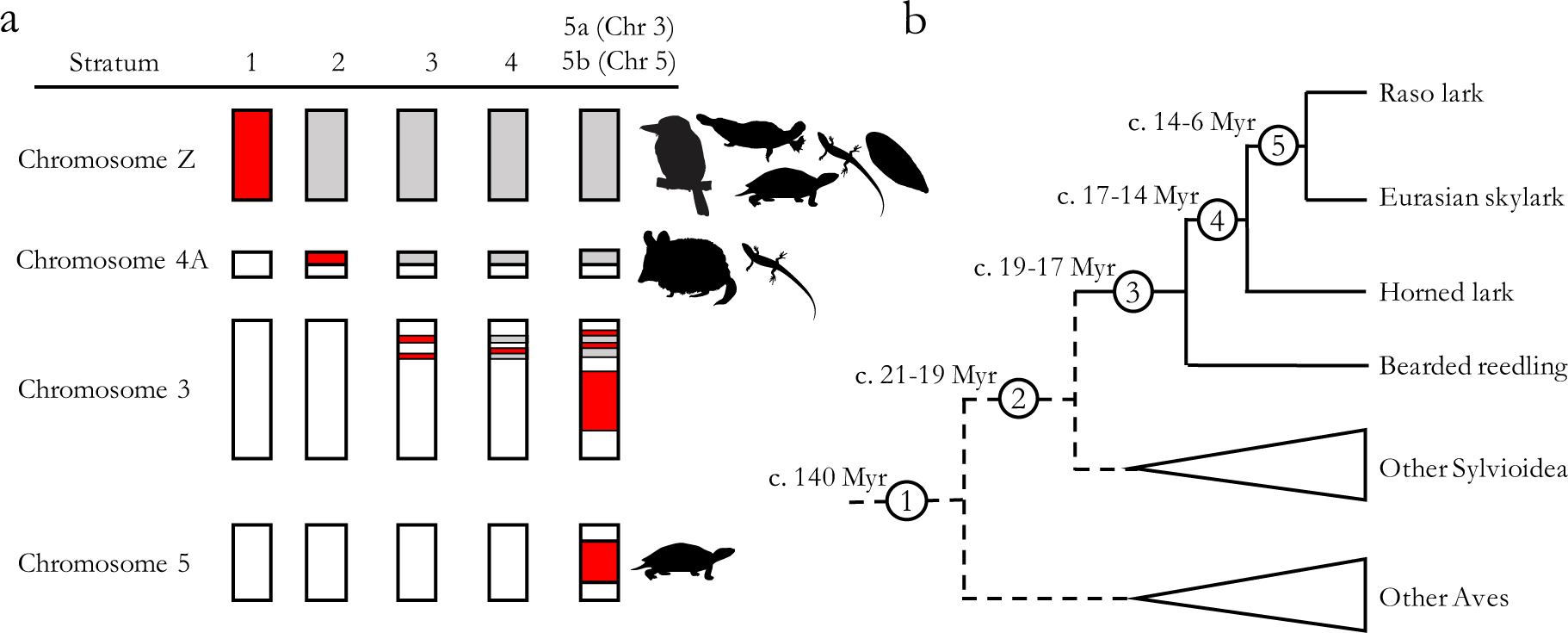
a) Cartoon representation of the evolutionary strata in the most parsimonious order of appearance (starting with stratum 1 and ending with stratum 5a and 5b). White colour represents autosomal regions and red colour marks the stratum being added. Grey colour represents sex-linked regions that already exist at the timing of the new stratum. Silhouettes indicate vertebrate groups where homologs to these chromosomes act as sex chromosomes (see Discussion): chromosome Z acts as the sex chromosome in all birds (Smith et al. 2009) and its homolog acts as sex chromosome in a flatfish (*Cynoglossus semilaevis*; Chen et al. 2014), a turtle (*Staurotypus triporcatus*; Montiel et al. 2016), a gekko (*Gekko hokouensis*; Kawai et al. 2008) and in the platypus (*Ornithorhynchus anatinus*; Grützner et al. 2004); the homolog of chromosome 4A is the sex chromosome in a lizard (*Takydromus sexlineatus*; Rovatsos et al. 2016) and in all eutherian mammals (Ross et al. 2005); and the homolog of chromosome 5 acts as sex chromosome in two turtle groups (*Glyptemys* wood turtles and *Siebenrockiella* marsh turtles; Montiel et al. 2016). (b) A cladogram showing when in time (Myr) the evolutionary strata were formed. The dating of stratum 1 is based on analyses of gametologous gene pairs in Cortez et al. (2014), stratum 2 and 3 on speciation estimates in Moyle et al. (2016), and stratum 4 and 5 (5a and 5b) on speciation estimates in Alström et al. (2013).

### 3.2 Analysis of divergence between gametologous genes

We analysed the Z–W divergence from the gametologous genes located on the different strata across the sex chromosome in the Raso lark and the Eurasian skylark, i.e. the species with all five evolutionary strata. We analysed the two different parts of stratum 5 on chromosome 3 and 5 separately in these analyses (i.e., stratum 5a and 5b, respectively). The degree of synonymous substitutions (median dS) between the Z and W gametologs ranged from 0.22 at stratum 1 to 0.02 at stratum 5a and 5b in the Raso lark, and from 0.21 at stratum 1 to 0.02 at stratum 5a and 5b in the Eurasian skylark (Figure 3, Suppl. Table S5). In the Raso lark, the log_2_-transformed dS values differed significantly between all strata except between stratum 2 and stratum 3, and between stratum 5a and 5b (Kruskal-Wallis rank sum test: chi-squared = 154.0, df = 4, p < 0.001; for pairwise comparison values see Suppl. Table S6). In the Eurasian skylark, the log_2_-transformed dS differed significantly between all strata, expect between stratum 5a and 5b (Kruskal-Wallis rank sum test: chi-squared = 143.0, df = 4, p < 0.001; pairwise comparison values in Suppl. Table S6). The degree of non-synonymous substitutions (dN) between the Z and W gametologs showed similar patterns, but less pronounced (median dN ranged from 0.03 at stratum 1 to 0.002 at stratum 5a in the Raso lark, and from 0.03 at stratum 1 to 0.002 at stratum 5a in the Eurasian skylark; Figure 3, Suppl. Table S5). The dN values were significantly different between stratum 5a and all other strata, and between 5b and all other strata in both species, and between stratum 1 and stratum 4 in the Raso lark (Raso lark: chi-squared = 88.7, df = 4, p < 0.001; Eurasian skylark: chi-squared = 143.0, df = 4, p < 0.001; for pairwise comparison values see Suppl. Table S6). The dN/dS values were lowest at stratum 1 in both species (0.082 in the Eurasian skylark and 0.097 in the Raso lark) and highest at stratum 4 (0.196 in the Eurasian skylark and 0.294 in the Raso lark). The dN/dS values in stratum 4 differed significantly from strata 1, 2 and 5a, and stratum 5a differed significantly from stratum 5b, in both species (Raso lark: chi-squared = 17.7, df = 4, p = 0.001; Eurasian skylark: chi-squared = 14.0, df = 4, p = 0.007; pairwise comparisons in Suppl. Table S6).

**Figure 3.**
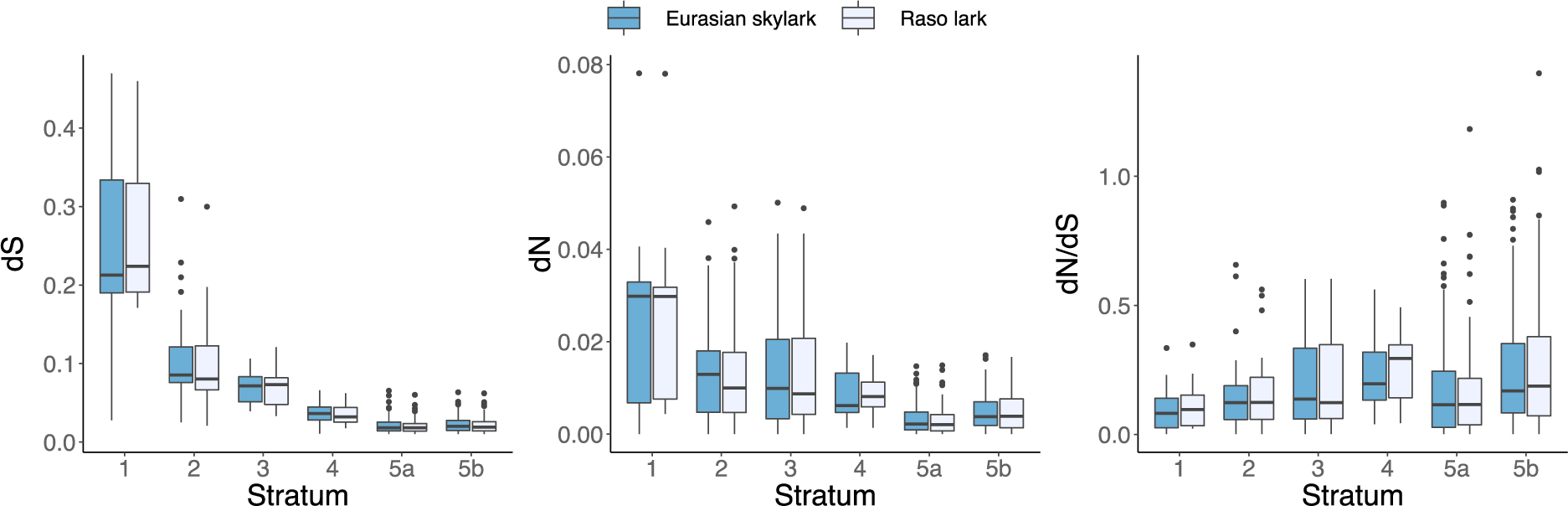
Z-to-W substitution rates for gametologous genes positioned within the different evolutionary strata in the Eurasian skylark (dark blue) and the Raso lark (light blue). (a) Median dS values, (b) median dN values, and (c) median dN/dS values are marked by the black line in each box, and the upper and lower hinges correspond to the first and third quarantiles (the 25 and 75 percentiles). The whiskers extend to no more than 1.5× the interquartile range from each hinge. See Suppl. Table S5 for details and main text for results from analyses of variances between strata based on log2-transformed values.

In both Raso larks and Eurasian skylarks, the dS values correlated significantly with the relative age of the strata (stratum 1 being oldest and strata 5a and 5b being youngest) (Ordered-heterogeneity test: r_s_P_c_ = 0.99, k = 6, p < 0.001 for both species). This was also true for the dN values (r_s_P_c_ = 0.99, k = 6, p < 0.001 for both species). However, the dN/dS values were not significantly associated with the age of the strata (Raso lark: r_s_P_c_ = 0.41, k = 6, p ∼ 0.11; Eurasian skylark: r_s_P_c_ = 0.46, k = 6, p ∼ 0.11).

### 3.3 Gene enrichment analysis

Of the 323 genes with sex-related GO term names (see Methods section), 16 genes were located on the ancestral avian sex chromosome Z (stratum 1), 5 within the sex-linked part of chromosome 4A (stratum 2), 33 within the sex-linked region of chromosome 3 (strata 3, 4 and 5a) and 5 within the sex-linked part of chromosome 5 (stratum 5b) (Figure 4a; Suppl. Table S7). The different strata on chromosome 3 hold 5 (stratum 3), 0 (stratum 4) and 28 (stratum 5a) genes, respectively (Figure 4b; Suppl. Table S7). This means that 59 genes with these hypothesised sex-related functions are sex-linked in the Raso lark and in the Eurasian skylark, i.e. the species with all strata, compared to 16 in species only having the ancestral avian sex chromosome.

Binomial tests showed that the sex-linked part of chromosome 3 (strata 3, 4 and 5a) had significantly more sex-related genes than would be expected if the genes were randomly distributed over the genome (33 observed genes compared to 19.7 expected genes; Binomial test: adjusted p = 0.019; Figure 4a; Suppl. Table S7). The sex-linked regions on chromosomes Z, 4A and 5 showed no statistical difference between observed and expected number of genes (NS; Suppl. Table S7). When analysing each stratum separately, stratum 5a was also significantly enriched for sex-related genes (28 observed genes compared to 16.7 expected genes; adjusted p = 0.047; Figure 4b; Suppl. Table S7), while the other strata were not significantly enriched (NS; Suppl. Table S7). Information about the analysed genes (chromosomal positions, GO terms and gene ID) are provided in Supplementary Document S1.

**Figure 4.**
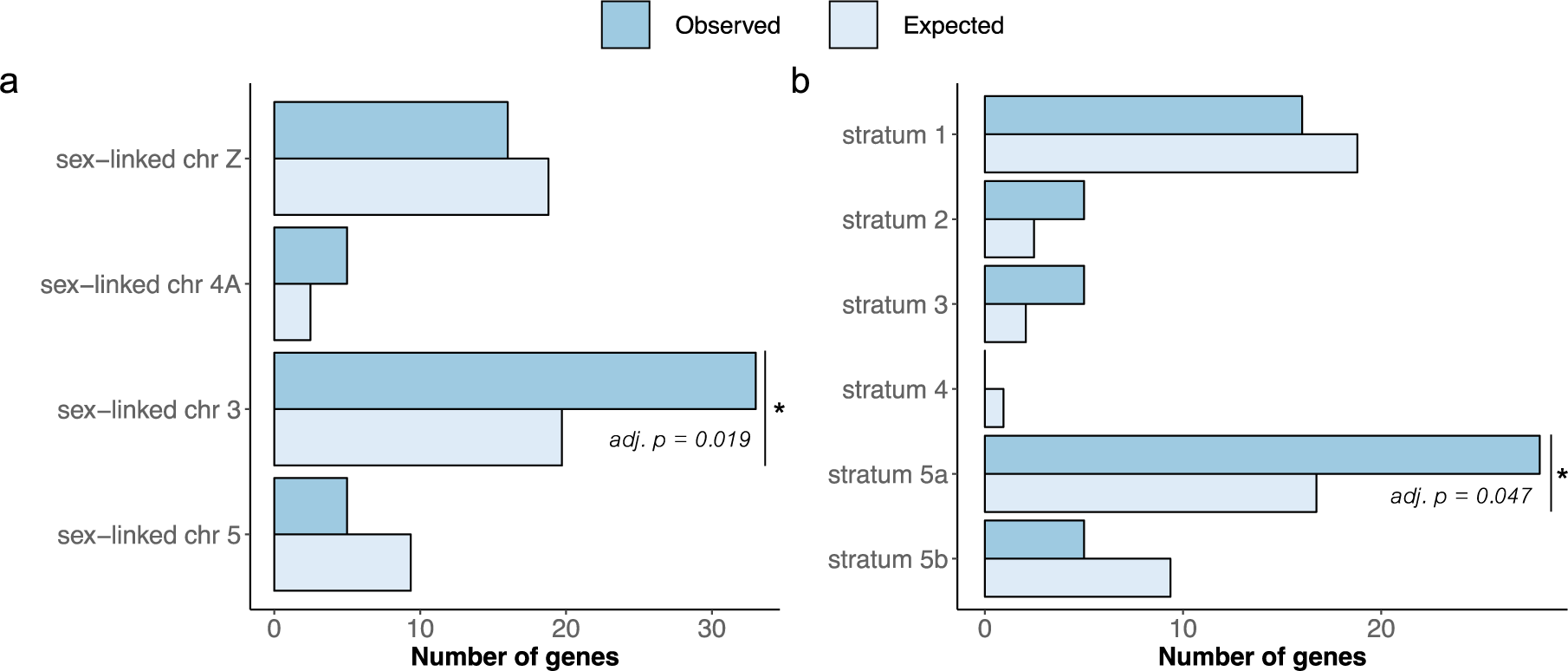
Observed and expected number of sex-related genes for (a) the sex-linked region of each chromosome and (b) each stratum. Binomial tests (Suppl. Table S7) showed that the sex-linked region on chromosome 3 had significantly more sex-related genes than would be expected if the genes were randomly distributed throughout the genome (adjusted p = 0.019), and that this enrichment on chromosome 3 was mainly due to a high number of sex-related genes on stratum 5a (adjusted p = 0.047).

## 4. DISCUSSION

By using comparative genomics in larks and their closest relative, the bearded reedling, we have identified the precise genomic regions that have evolved sex-linkage across a group of birds with multiple autosome– sex chromosome fusions. The extent of sex-linkage varied between species, and for two of them (Raso lark and Eurasian skylark in the genus *Alauda*), we detected the largest known avian sex chromosome (195.3 Mbp) and show that it originates from fusions between (parts of) four avian chromosomes (Z, 3, 4A and 5). We found evidence of five evolutionary strata where recombination has been suppressed at different time points; from c. 140 Myr ago (stratum 1, the sex chromosome common to all birds) to c. 14-6 Myr ago (stratum 5a and 5b, the layer unique to *Alauda*; the strata and their age estimates are summarised in Figure 2). We further show that the substitution rates between Z and W gametologs increase with increasing age of the strata. This gives support for a key prediction from sex chromosome theory: continuous differentiation of non-recombining chromosomes (Rice 1994; Lahn & Page 1999).

With the available genomic data, we are not able to determine the chromosomal structure of this neo-sex chromosome. However, the 195.3 Mbp sex-linked region in the two *Alauda* study species (Raso lark and Eurasian skylark) constitutes c. 16.3% of their expected genome size (1.2 Gbp), which in turn corresponds well to the genomic proportion of the sex chromosomes in the karyotypes of the bimaculated lark (i.e., c. 15-20%; Bulatova 1981). The bimaculated lark karyotypes further show that the Z and W chromosomes are of similar size and drastically enlarged compared to the usual bird karyotype (Bulatova 1981). Therefore, we believe that the additional chromosomes that are sex-linked in larks have fused to the Z as well as the W chromosome. Moreover, the karyotype of a male horned lark (no female was karyotyped) indicates a Z chromosome of similar size to that of the bimaculated lark (Bulatova 1981). Therefore, it is possible that large parts of chromosome 3 and 5 are fused to the Z chromosome also in the horned lark, but as our results show are still recombining in that species (with the exception of strata 3 and 4 on chromosome 3 that are non-recombining). This reasoning is complemented by the results in E. Dierickx et al. (unpublished manuscript) showing that different species and subspecies of *Alauda* (the oriental skylark *A. gulgula*, Eurasian skylark sspp. [possibly *A. a. arvensis*/*dulcivox*/*intermedia*], and Raso lark), and a species of the genus *Gallerida* (the crested lark *G. cristata*) – the sister genus of *Alauda* – have suppressed recombination on chromosome 5 (stratum 5b) but show recombination suppression to different degrees on chromosome 3 (stratum 5a).

A long-standing hypothesis in sex chromosome research posits that the evolutionary dynamics of sex chromosomes are governed by selection acting on linked genes, in particular on sex-determining and sexually antagonistic genes (Haldane 1922; Fisher 1931; Lewis & John 1968; D. Charlesworth & B. Charlesworth 1980). This hypothesis states that chromosomes that harbour such genes should be over-represented in sex chromosome formations, transitions and turnovers (Bachtrog et al. 2011; Ross et al. 2009; O’Meally et al. 2012). In support of this hypothesis, we found that the sex-linked region of chromosome 3, and one of the strata on that chromosome (stratum 5a), were significantly enriched for genes with sex-related functions (adjusted p = 0.019 and 0.047, respectively). Among these genes is *ESR1* (*Estrogen receptor 1*), which has been suggested to be involved in sex reversal in the American alligator (*Alligator mississippiensis*; Kohno et al. 2015). The other chromosomes (and strata) involved in the extensive neo-sex chromosome formation in larks were not significant enriched for sex-related genes. However, all of them have independently been recruited as sex chromosomes in other vertebrate lineages. The Z chromosome – the sex chromosome in all birds, containing the putative sex determining gene *DMRT1* (*Doublesex and mab-3 related transcription factor 1*; Smith et al. 2009) – has independently been recruited as a sex chromosome in a flatfish (the half-smooth tongue sole *Cynoglossus semilaevis*; Chen et al. 2014), a turtle (the Mexican musk turtle *Staurotypus triporcatus*; Montiel et al. 2016), a gecko (the Kwangsi gecko *Gekko hokouensis;* Kawai et al. 2008), and in the platypus (*Ornithorhynchus anatinus*; Grützner et al. 2004). Chromosome 4A is homologous to the sex chromosome in a lizard (the Asian grass lizard *Takydromus sexlineatus*; Rovatsos et al. 2016) and in all eutherian mammals (Ross et al. 2005), and contains several interesting genes, including *AR* (*Androgen receptor*) and *SOX3* (*SRY-related HMG-box 3*) (note, however, that *SOX3* in not sex-linked in Sylvioidea; Pala, Naurin, et al. 2012; this study). Finally, the homolog to chromosome 5 acts as sex chromosome in two turtle groups (*Glyptemys insculpta* and *Siebenrockiella crassicollis*; Montiel et al. 2016). An alternative hypothesis is that fusions may become fixed as a consequence of non-selective processes (see e.g. Pennell et al. 2015). However, we have presented several strands of evidence – the enrichment of genes with sex-specific function and the repeated homology to other vertebrate sex chromosomes – to support the hypothesis that the multiple fusion events between chromosome Z, 4A, 3 and 5, that have formed these extraordinary neo-sex chromosomes in larks and in the bearded reedling, have been driven by selective processes acting on their gene content. This adds to the accumulating evidences that specific chromosomes are non-randomly recruited as sex chromosomes in vertebrates (O’Meally et al. 2012; Ezaz et al. 2006; Ezaz et al. 2016).

Multiple autosome–sex chromosome fusions have also occurred in other clades, including gazelles and snakes (Vassart et al. 1995; Pokorná et al. 2014). In snakes, all six neo-sex chromosome systems identified so far are from a single family, Elapidae (Pokorná et al. 2014). These patterns, now also including those found in the larks, hint at a phylogenetic component in neo-sex chromosome formation; that once an autosome–sex chromosome fusion has occurred, additional ones may be expected (Pokorná et al. 2014). If the fusions are being selected for because of any advantageous linkage that they may create, then previously fused regions (with already suppressed recombination) provide more opportunities for beneficial linkage to be formed between the sex-linked genes and new additional genomic regions.

## Acknowledgements

We wish to thank O. Berglund, P. Zehtindjiev and S. Bensch for providing DNA of our study species. Sequencing was performed by the SNP&SEQ Technology Platform in Uppsala, which is part of the National Genomics Infrastructure (NGI) Sweden and Science for Life Laboratory, supported by the Swedish Research Council and the Knut and Alice Wallenberg Foundation. Bioinformatics analyses were performed on resources provided by the Swedish National Infrastructure for Computing (SNIC) at Uppsala Multidisciplinary Center for Advanced Computational Science (UPPMAX). The research was funded by research grants from the Swedish Research Council (to BH: 621-2014-5222 and 621-2016-689), the Royal Physiological Society in Lund (the Nilsson-Ehle Foundation), the Erik Philip-Sörensens Foundation, the Stiftelsen Olle Engkvist and the Wenner-Gren Foundations (to SP).

## Supplementary Material and Data Availability

Suppl. Figure S1 and Suppl. Tables S1-7 are provided together as a separate file. Suppl. Document S1 is provided as a separate file. In-house scripts are given in Suppl. Code S1 (available upon request). Illumina HiSeqX raw reads (150 bp, paired-end) will been deposited at NCBI sequence read archives upon acceptance.

## References

Alström, P. et al., 2013. Multilocus phylogeny of the avian family Alaudidae (larks) reveals complex morphological evolution, non-monophyletic genera and hidden species diversity. Molecular Phylogenetics and Evolution, 69(3), pp.1043–1056.

Bachtrog, D. et al., 2011. Are all sex chromosomes created equal? Trends in genetics: TIG, 27(9), pp.350–357. Available at: http://dx.doi.org/10.1016/j.tig.2011.05.005.

Bankevich, A. et al., 2012. SPAdes: a new genome assembly algorithm and its applications to single-cell sequencing. Journal of computational biology: a journal of computational molecular cell biology, 19(5), pp.455–477.

Bolger, A.M., Lohse, M. & Usadel, B., 2014. Trimmomatic: a flexible trimmer for Illumina sequence data. Bioinformatics, 30(15), pp.2114–2120.

Brooke, M. de L. et al., 2010. Widespread translocation from autosomes to sex chromosomes preserves genetic variability in an endangered lark. Journal of Molecular Evolution, 70(3), pp.242–246.

Bulatova, N.S., 1981. A comparative karyological study of passerine bird.

Castresana, J., 2000. Selection of conserved blocks from multiple alignments for their use in phylogenetic analysis. Molecular Biology and Evolution, 17(4), pp.540–552.

Charlesworth, D. & Charlesworth, B., 1980. Sex differences in fitness and selection for centric fusions between sex-chromosomes and autosomes. Genetical research, 35(2), pp.205–214. Available at: http://www.ncbi.nlm.nih.gov/pubmed/6930353.

Chen, S. et al., 2014. Whole-genome sequence of a flatfish provides insights into ZW sex chromosome evolution and adaptation to a benthic lifestyle. Nature Genetics, 46(3), pp.253–260. Available at: http://www.ncbi.nlm.nih.gov/pubmed/24487278.

Cortez, D. et al., 2014. Origins and functional evolution of Y chromosomes across mammals. Nature, 508(7497), pp.488–493. Available at: http://www.ncbi.nlm.nih.gov/pubmed/24759410.

Cunningham, F. et al., 2015. Ensembl 2015. Nucleic Acids Research, 43, pp.D662–9. Available at: http://www.pubmedcentral.nih.gov/articlerender.fcgi?artid=PMC4383879.

Danecek, P. et al., 2011. The variant call format and VCFtools. Bioinformatics, 27(15), pp.2156–2158.

Ellegren, H., 2010. Evolutionary stasis: the stable chromosomes of birds. Trends in Ecology & Evolution, 25(5), pp.283–291.

Ezaz, T. et al., 2006. Relationships between Vertebrate ZW and XY Sex Chromosome Systems. Current biology: CB, 16(17), pp.R736–R743.

Ezaz, T., Srikulnath, K. & Graves, J.A.M., 2016. Origin of Amniote Sex Chromosomes: An Ancestral Super-Sex Chromosome, or Common Requirements? Journal of Heredity, 108(1), pp.94–105.

Fisher, R.A., 1931. THE EVOLUTION OF DOMINANCE. Biological Reviews, 6(4), pp.345–368. Available at: http://doi.wiley.com/10.1111/j.1469-185X.1931.tb01030.x.

Garrison, E. & Marth, G., 2012. Haplotype-based variant detection from short-read sequencing. arxiv.org.

Grabherr, M.G. et al., 2010. Genome-wide synteny through highly sensitive sequence alignment: Satsuma. Bioinformatics, 26(9), pp.1145–1151.

Grützner, F. et al., 2004. In the platypus a meiotic chain of ten sex chromosomes shares genes with the bird Z and mammal X chromosomes. Nature, 432(7019), pp.913–917.

Gurevich, A. et al., 2013. QUAST: quality assessment tool for genome assemblies. Bioinformatics, 29(8), pp.1072–1075.

Haldane, J.B.S., 1922. Sex ratio and unisexual sterility in hybrid animals. Journal of Genetics, 12(2), pp.101–109.

Kawai, A. et al., 2008. The ZW sex chromosomes of *Gekko hokouensis* (Gekkonidae, Squamata) represent highly conserved homology with those of avian species. Chromosoma, 118(1), pp.43–51.

Kielbasa, S.M. et al., 2011. Adaptive seeds tame genomic sequence comparison. Genome Research, 21(3), pp.487–493.

Kitano, J. & Peichel, C.L., 2011. Turnover of sex chromosomes and speciation in fishes. Environmental Biology of Fishes, 94(3), pp.549–558.

Kitano, J. et al., 2009. A role for a neo-sex chromosome in stickleback speciation. Nature, 461(7267), pp.1079–1083.

Kohno, S. et al., 2015. Estrogen Receptor 1 (ESR1; ERα), not ESR2 (ERβ), Modulates Estrogen-Induced Sex Reversal in the American Alligator, a Species With Temperature-Dependent Sex Determination. Endocrinology, 156(5), pp.1887–1899.

Lahn, B.T. & Page, D.C., 1999. Four Evolutionary Strata on the Human X Chromosome. Science, 286(5441), pp.964–967.

Lewis, K.R. & John, B., 1968. The Chromosomal Basis of Sex Determination. In International Review of Cytology. Elsevier, pp. 277–379.

Li, H. & Durbin, R., 2009. Fast and accurate short read alignment with Burrows–Wheeler transform. Bioinformatics, 25(14), pp.1754–1760.

Li, H. et al., 2009. The Sequence Alignment/Map format and SAMtools. Bioinformatics, 25(16), pp.2078– 2079.

Loytynoja, A., 2014. Phylogeny-aware alignment with PRANK. Methods in molecular biology (Clifton, N.J.), 1079(Chapter 10), pp.155–170.

Montiel, E.E. et al., 2016. Cytogenetic Insights into the Evolution of Chromosomes and Sex Determination Reveal Striking Homology of Turtle Sex Chromosomes to Amphibian Autosomes. Cytogenetic and Genome Research, 148(4), pp.292–304.

Moyle, R.G. et al., 2016. Tectonic collision and uplift of Wallacea triggered the global songbird radiation. 7, 12709 EP –.

O’Meally, D. et al., 2012. Are some chromosomes particularly good at sex? Insights from amniotes. Chromosome Research, 20(1), pp.7–19.

Pala, I., Hasselquist, D., et al., 2012. Patterns of molecular evolution of an avian neo-sex chromosome. Molecular Biology and Evolution, 29(12), pp.3741–3754.

Pala, I., Naurin, S., et al., 2012. Evidence of a neo-sex chromosome in birds. Heredity, 108(3), pp.264–272. Available at: http://dx.doi.org/10.1038/hdy.2011.70.

Pennell, M. W. et al., 2015. Y Fuse? Sex Chromosome Fusions in Fishes and Reptiles. PLOS Genetics, 11(5), e1005237. Available at: http://doi.org/10.1371/journal.pgen.1005237

Pokorná, M., Altmanová, M. & Kratochvíl, L., 2014. Multiple sex chromosomes in the light of female meiotic drive in amniote vertebrates. Chromosome Research, 22(1), pp.35–44.

Quinlan, A.R. & Hall, I.M., 2010. BEDTools: a flexible suite of utilities for comparing genomic features. Bioinformatics, 26(6), pp.841–842.

Rice, W.R., 1994. Degeneration of a nonrecombining chromosome. Science, 263(5144), pp.230–232.

Rice, W.R. & Gaines, S.D., 1994. Extending nondirectional heterogeneity tests to evaluate simply ordered alternative hypotheses. Proceedings of the National Academy of Sciences, 91(1), pp.225–226.

Rovatsos, M. et al., 2016. Mammalian X homolog acts as sex chromosome in lacertid lizards. Heredity, 117(1), 8–13. Available at: http://doi.org/10.1038/hdy.2016.18

Ross, J.A. et al., 2009. Turnover of Sex Chromosomes in the Stickleback Fishes (Gasterosteidae). PLOS Genetics, 5(2), p.e1000391.

Ross, M.T et al. 2005. The DNA sequence of the human X chromosome. Nature, 434(7031), 325–337. http://doi.org/10.1038/nature03440

Sambrock, J. & W Russel, D, 2001. Molecular Cloning: A Laboratory Manual (3rd edition). Cold Spring Harbor Laboratory Press.

Sigeman, H. et al., 2018. Insights into Avian Incomplete Dosage Compensation: Sex-Biased Gene Expression Coevolves with Sex Chromosome Degeneration in the Common Whitethroat. Genes, 9(8), p.373.

Smith, C.A. et al., 2009. The avian Z-linked gene DMRT1 is required for male sex determination in the chicken. Nature, 461(7261), pp.267–271.

Tange, O., 2018. GNU Parallel 2018, Lulu.com.

Vassart, M., Séguéla, A. & Hayes, H., 1995. Chromosomal evolution in gazelles. Journal of Heredity, 86(3), pp.216–227.

Warren, W.C. et al., 2010. The genome of a songbird. Nature, 464(7289), pp.757–762.

Yang, Z., 2007. PAML 4: Phylogenetic Analysis by Maximum Likelihood. Molecular Biology and Evolution, 24(8), pp.1586–1591.

Yoshida, K. et al., 2014. Sex Chromosome Turnover Contributes to Genomic Divergence between Incipient Stickleback Species J. Zhang, ed. PLOS Genetics, 10(3), p.e1004223.

Zamani, N. et al., 2014. A universal genomic coordinate translator for comparative genomics. BMC Bioinformatics, 15(1), p.227.

Zhou, Q. et al., 2014. Complex evolutionary trajectories of sex chromosomes across bird taxa. Science, 346(6215), pp.1246338–1246338. Available at: http://www.sciencemag.org/cgi/doi/10.1126/science.1246338.

