## Supplementary Figures and Tables for "Genomics of expanded avian sex chromosomes shows that certain chromosomes are predisposed towards sex-linkage in vertebrates"

### Supplementary Materials:

#### Supplementary Figures

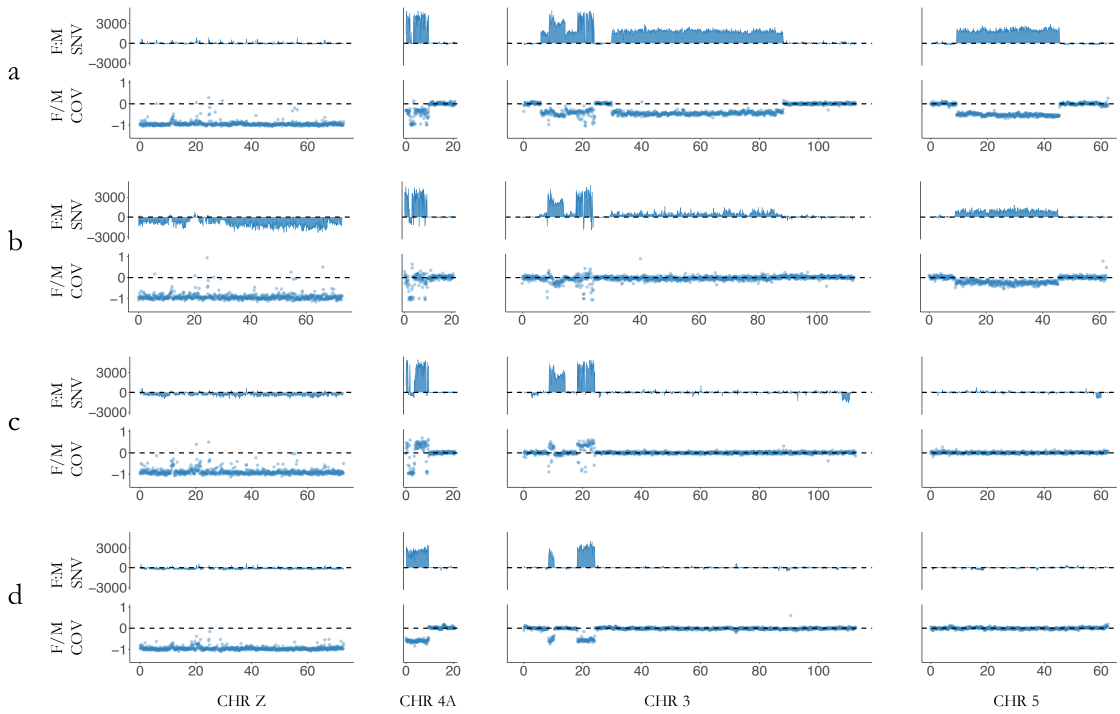


**Suppl. Figure S1.** Mean female-to-male difference in number of private single nucleotide variants (SNVs) and female-to-male coverage ratios across 0.1 Mbp windows for the four chromosomes (Z, 4A, 3 and 5) with genomic patterns of sex-linkage. a) Raso lark, b) Eurasian skylark, c) horned lark and d) bearded reedling.

### Supplementary Tables

**Suppl. Table S1.** Genome assembly statistics computed using the software Quast version 4.5.4 [1].

| Assembly | Raso lark | Horned lark | Eurasian skylark | Bearded reedling |
|---|---|---|---|---|
| *N* contigs (>= 0 bp) | 28304 | 109088 | 256822 | 36455 |
| *N* contigs (>= 1000 bp) | 28304 | 109088 | 256822 | 36455 |
| *N* contigs (>= 5000 bp) | 17977 | 51289 | 79725 | 24408 |
| *N* contigs (>= 10000 bp) | 14382 | 32587 | 31198 | 19461 |
| *N* contigs (>= 25000 bp) | 9767 | 10545 | 3546 | 12025 |
| *N* contigs (>= 50000 bp) | 6145 | 2805 | 609 | 6436 |
| Total length (>= 0 bp) | 1003896371 | 1086584534 | 1276061662 | 1020354129 |
| Total length (>= 1000 bp) | 1003896371 | 1086584534 | 1276061662 | 1020354129 |
| Total length (>= 5000 bp) | 979649475 | 962416262 | 887743951 | 991752640 |
| Total length (>= 10000 bp) | 953912007 | 827105499 | 543408884 | 955927387 |
| Total length (>= 25000 bp) | 878299503 | 479556955 | 145311035 | 833768162 |
| Total length (>= 50000 bp) | 748288932 | 216102095 | 49632478 | 634399538 |
| *N* contigs | 28304 | 109088 | 256822 | 36455 |
| Largest contig (bp) | 851956 | 447328 | 346983 | 534950 |
| Total length (bp) | 1003896371 | 1086584534 | 1276061662 | 1020354129 |
| GC (%) | 42.31 | 42.17 | 42.59 | 42.14 |
| N50 | 103210 | 21578 | 8477 | 68188 |
| N75 | 49107 | 10418 | 4130 | 32980 |
| L50 | 2726 | 13291 | 41480 | 4314 |
| L75 | 6239 | 31395 | 94969 | 9642 |
| # N's per 100 kbp | 0.87 | 16.02 | 23.63 | 0.85 |

[1] Gurevich, A., Saveliev, V., Vyahhi, N. & Tesler, G. QUAST: quality assessment tool for genome assemblies. *Bioinformatics* **29,** 1072–1075 (2013).

**Suppl. Table S2.** Alignment statistics for the female and male sample from the four studied species (see Methods section 2.2 for details). No. read pairs refer to the number of read pairs remaining after quality trimming. The number of properly aligning reads was calculated with samtools v1.7 flagstat [2].

| | | | | | Properly paired mapped reads (after deduplication) | |
|---|---|---|---|---|---|---|
| Sample | Species | Reference genome | Sex | No. read pairs | All mismatches allowed | <=2 mismatches |
| AlaRaz_f | Raso lark | Raso lark | female | 101453455 | 174881744 | 161376774 |
| AlaRaz_m | Raso lark | Raso lark | male | 87822864 | 152776522 | 147354471 |
| AlaArv_f | Eurasian skylark | Raso lark | female | 151270768 | 224364208 | 111786956 |
| AlaArv_m | Eurasian skylark | Raso lark | male | 147067636 | 227283590 | 114762381 |
| EreAlp_f | Horned lark | Raso lark | female | 130242267 | 180857630 | 69507055 |

| | | | | | | |
|---|---|---|---|---|---|---|
| EreAlp_m | Horned lark | Raso lark | male | 202099679 | 275199656 | 104740879 |
| PanBia_f | Bearded reedling | Bearded reedling | female | 98173644 | 138219068 | 125403224 |
| PanBia_m | Bearded reedling | Bearded reedling | male | 163911536 | 230747108 | 218069639 |

[2] Li, H. *et al.* The Sequence Alignment/Map format and SAMtools. *Bioinformatics* **25,** 2078–2079 (2009).

**Suppl. Table S3.** Gametolog extraction criteria. Z and W gametolog sequences were extracted based on the genotype distributions between the male and female sample.

| **Variant type** | **Nr of mutations** | **Male** | **Female** | **Extracted Z allele** | **Extracted W allele** |
|---|---|---|---|---|---|
| biallelic | 0 | 0/0 | 0/0 | 0 | 0 |
| | 1 | 0/1 | 0/0 | N | N |
| | 1 | 0/0 | 0/1 | 0 | 1 |
| | 2 | 0/1 | 0/1 | N | N |
| | 3 | 1/1 | 0/1 | 1 | 0 |
| | 3 | 0/1 | 1/1 | N | N |
| | 4 | 1/1 | 1/1 | 1 | 1 |
| triallelic | 1 | 1/2 | 1/1 | N | 1 |
| | 1 | 1/1 | 1/2 | 1 | 2 |
| | 2 | 1/2 | 1/2 | N | N |
| | 3 | 2/2 | 1/2 | 2 | 1 |
| | 3 | 1/2 | 2/2 | N | N |
| | 4 | 2/2 | 2/2 | 2 | 2 |

0/0 = no variation in relation to reference genome

0/1 = heterozygote genotype in relation to reference genome

1/1 = homozygote genotype with one allele not found in reference genome

1/2 = heterozygote genotype with two different variants not found in reference genome

2/2 = homozygote genotype not found in reference genome

**Suppl. Table S4.** The mean female-to-male coverage ratio (cov) and difference in number of female-specific mutations and male-specific mutations (snp) of all 1 Mbp windows per chromosome.

| Chromosome | Raso lark | | Eurasian skylark | | Horned lark | | Bearded reedling | |
|---|---|---|---|---|---|---|---|---|
| | coverage | snp | coverage | snp | coverage | snp | coverage | snp |
| 1 | 1.03 | -55.76 | 1.01 | 59.39 | 1.06 | -428.35 | 0.99 | -90.65 |
| 1A | 1.04 | -453.08 | 1.02 | 174.25 | 1.07 | 265.73 | 1.00 | -193.37 |
| 1B | 0.98 | 130.50 | 1.03 | 278.00 | 1.07 | -83.00 | 1.04 | 37.00 |
| 2 | 1.03 | 129.38 | 1.00 | 91.55 | 1.06 | 46.71 | 0.99 | -30.68 |

| | | | | | | | | |
|---|---|---|---|---|---|---|---|---|
| 3 | 0.84 | 13459.84 | 0.98 | 4958.29 | 1.07 | 2775.37 | 0.97 | 1898.89 |
| 4 | 1.02 | 218.61 | 1.00 | -121.41 | 1.05 | 503.89 | 0.99 | -121.34 |
| 4A | 0.90 | 13033.77 | 0.96 | 9762.91 | 1.09 | 10978.41 | 0.85 | 10497.82 |
| 5 | 0.84 | 11011.22 | 0.91 | 5102.22 | 1.06 | -90.25 | 0.99 | -122.97 |
| 6 | 1.02 | -64.81 | 1.17 | 52.16 | 1.08 | 43.51 | 1.00 | 78.08 |
| 7 | 1.03 | 119.59 | 1.00 | -65.32 | 1.07 | -11.24 | 1.00 | -64.37 |
| 8 | 1.02 | -309.62 | 1.00 | 3.86 | 1.05 | -6.00 | 1.00 | -62.31 |
| 9 | 1.03 | -727.96 | 1.01 | 409.54 | 1.05 | -5353.43 | 1.00 | 21.68 |
| 10 | 1.02 | 950.95 | 1.00 | 55.23 | 1.06 | -1323.82 | 1.01 | -171.36 |
| 11 | 1.03 | 637.95 | 1.01 | 47.00 | 1.06 | -18.09 | 1.01 | -275.05 |
| 12 | 1.03 | 130.96 | 1.01 | -15.91 | 1.06 | 74.39 | 1.01 | -9.70 |
| 13 | 1.02 | 191.17 | 1.00 | -6.50 | 1.06 | -120.17 | 1.01 | -367.50 |
| 14 | 1.02 | 107.88 | 1.01 | 132.82 | 1.06 | 99.76 | 1.01 | 99.94 |
| 15 | 1.02 | -902.73 | 1.01 | 9.80 | 1.06 | -8.47 | 1.02 | -188.53 |
| 17 | 1.02 | -112.08 | 1.00 | 80.69 | 1.06 | 140.23 | 1.02 | -448.00 |
| 18 | 1.02 | -666.17 | 1.00 | -43.50 | 1.06 | -71.42 | 1.01 | 467.58 |
| 19 | 1.03 | 102.77 | 1.01 | 97.54 | 1.07 | 64.31 | 1.03 | 38.54 |
| 20 | 1.02 | -137.18 | 1.00 | 18.59 | 1.07 | -19.94 | 1.02 | -65.41 |
| 21 | 1.02 | -880.43 | 1.03 | 52.14 | 1.07 | -1986.71 | 1.02 | -563.29 |
| 22 | 1.03 | 246.75 | 1.00 | 27.75 | 1.06 | 328.00 | 1.01 | 716.75 |
| 23 | 1.02 | -780.00 | 1.01 | 76.00 | 1.07 | -99.57 | 1.03 | -20.29 |
| 24 | 1.03 | -355.78 | 1.00 | 57.00 | 1.07 | -78.22 | 1.02 | 285.00 |
| 25 | 1.03 | -99.50 | 1.05 | -681.50 | 1.08 | 231.00 | 1.04 | 90.50 |
| 26 | 1.02 | -198.17 | 1.02 | -1245.50 | 1.06 | -95.50 | 1.03 | 103.00 |
| 27 | 1.02 | 336.33 | 1.00 | 93.33 | 1.07 | -228.83 | 1.03 | 160.33 |
| 28 | 1.03 | -52.50 | 1.05 | 187.83 | 1.35 | 134.33 | 1.05 | -35.83 |
| Z | 0.53 | -174.12 | 0.54 | -9719.91 | 0.57 | -3682.34 | 0.51 | -1570.14 |

**Suppl. Table S5.** Nucleotide substitution values for gametologous (Z-W) gene pairs from Raso lark and Eurasian skylark for each sex chromosome strata.

| Stratum | Chromosome | | No. genes | Median dS | Median dN | Median dN/dS |
|---|---|---|---|---|---|---|
| 1 | Z | Raso lark | 9 | 0.224 | 0.030 | 0.097 |
| | | Eurasian skylark | 10 | 0.213 | 0.030 | 0.082 |
| 2 | 4A | Raso lark | 33 | 0.080 | 0.010 | 0.124 |
| | | Eurasian skylark | 32 | 0.085 | 0.013 | 0.123 |
| 3 | 3 | Raso lark | 23 | 0.073 | 0.009 | 0.123 |
| | | Eurasian skylark | 23 | 0.072 | 0.010 | 0.137 |

| | | | | | | |
|---|---|---|---|---|---|---|
| 4 | 3 | Raso lark | 18 | 0.032 | 0.008 | 0.294 |
| | | Eurasian skylark | 18 | 0.036 | 0.006 | 0.196 |
| 5a | 3 | Raso lark | 168 | 0.018 | 0.002 | 0.116 |
| | | Eurasian skylark | 186 | 0.018 | 0.002 | 0.115 |
| 5b | 5 | Raso lark | 164 | 0.019 | 0.004 | 0.187 |
| | | Eurasian skylark | 169 | 0.020 | 0.004 | 0.168 |

**Suppl. Table S6.** P values from statistical tests of nucleotide differences between pairs of evolutionary strata (see Methods).

| | | Stratum 1 | Stratum 2 | Stratum 3 | Stratum 4 | Stratum 5a |
|---|---|---|---|---|---|---|
| *Analysis of synonymous substitutions (dS)* | | | | | | |
| Eurasian skylark | Stratum 2 | < 0.001 | | | | |
| Raso lark | | < 0.001 | | | | |
| Eurasian skylark | Stratum 3 | < 0.001 | 0.016 | | | |
| Raso lark | | < 0.001 | 0.091 | | | |
| Eurasian skylark | Stratum 4 | < 0.001 | < 0.001 | < 0.001 | | |
| Raso lark | | < 0.001 | < 0.001 | < 0.001 | | |
| Eurasian skylark | Stratum 5a | < 0.001 | < 0.001 | < 0.001 | < 0.001 | |
| Raso lark | | < 0.001 | < 0.001 | < 0.001 | < 0.001 | |
| Eurasian skylark | Stratum 5b | < 0.001 | < 0.001 | < 0.001 | < 0.001 | 0.087 |
| Raso lark | | < 0.001 | < 0.001 | < 0.001 | < 0.001 | 0.185 |
| *Analysis of non-synonymous substitutions (dN)* | | | | | | |
| Eurasian skylark | Stratum 2 | 0.255 | | | | |
| Raso lark | | 0.088 | | | | |
| Eurasian skylark | Stratum 3 | 0.232 | 0.993 | | | |
| Raso lark | | 0.085 | 0.894 | | | |
| Eurasian skylark | Stratum 4 | 0.055 | 0.272 | 0.360 | | |
| Raso lark | | 0.038 | 0.292 | 0.421 | | |
| Eurasian skylark | Stratum 5a | < 0.001 | < 0.001 | < 0.001 | < 0.001 | |
| Raso lark | | < 0.001 | < 0.001 | < 0.001 | < 0.001 | |
| Eurasian skylark | Stratum 5b | 0.002 | < 0.001 | 0.000 | 0.002 | < 0.001 |
| Raso lark | | < 0.001 | < 0.001 | 0.000 | < 0.001 | < 0.001 |
| *Analysis of rate of evolution (dN/dS)* | | | | | | |
| Eurasian skylark | Stratum 2 | 0.402 | | | | |
| Raso lark | | 0.527 | | | | |
| Eurasian skylark | Stratum 3 | 0.271 | 0.488 | | | |
| Raso lark | | 0.306 | 0.527 | | | |
| Eurasian skylark | Stratum 4 | 0.040 | 0.043 | 0.402 | | |
| Raso lark | | 0.028 | 0.028 | 0.303 | | |

| | | | | | | |
|---|---|---|---|---|---|---|
| Eurasian skylark | Stratum 5a | 0.488 | 0.875 | 0.402 | 0.043 | |
| Raso lark | | 0.864 | 0.527 | 0.303 | 0.003 | |
| Eurasian skylark | Stratum 5b | 0.147 | 0.190 | 0.746 | 0.416 | 0.032 |
| Raso lark | | 0.303 | 0.303 | 0.678 | 0.341 | 0.002 |

**Suppl. Table S7.** Results from binominal tests for significant enrichment of genes involved in a range of sex-related functions (see Methods section 2.6 for details). "Sex-related genes within region" corresponds to the observed genes in Figure 5. Expected genes in Figure 5 was calculated as the total number of sex-related genes within the genome (n = 323) divided by "Proportion of genome". Genes mentioned in the Discussion section are marked in red.

| Chromosome | Region | Proportion of genome | | Sex-related genes | | P value | Adjusted p value |
|---|---|---|---|---|---|---|---|
| *Strata level analysis* | | Observed | Estimated (CI) | Genome-wide | Within region | | |
| Z | 1 | 0.058 | 0.05 (0.029-0.079) | 323 | 16[#] | 0.633 | 1 |
| 4A | 2 | 0.008 | 0.015 (0.005-0.036) | 323 | 5[$] | 0.105 | 0.419 |
| 3 | 3 | 0.006 | 0.015 (0.005-0.036) | 323 | 5[&] | 0.058 | 0.289 |
| 3 | 4 | 0.003 | 0 (0.000-0.011) | 323 | 0 | 1 | 1 |
| 3 | 5a | 0.052 | 0.087 (0.058-0.123) | 323 | 28€ | 0.008 | **0.047** |
| 5 | 5b | 0.029 | 0.015 (0.005-0.036) | 323 | 5[!] | 0.183 | 0.548 |
| *Total sex-linked region per chromosome analysis* | | | | | | | |
| Z | 1 | 0.058 | 0.05 (0.029-0.079) | 323 | 16 | 0.633 | 0.633 |
| 4A | 2 | 0.008 | 0.015 (0.005-0.036) | 323 | 5 | 0.105 | 0.314 |
| 3 | 3 + 4 + 5a | 0.061 | 0.1 (0.071-0.140) | 323 | 33 | 0.005 | **0.019** |
| 5 | 5b | 0.029 | 0.015 (0.005-0.036) | 323 | 5 | 0.183 | 0.365 |

*# B4GALT1, CRHBP, DDX4, DMRT1, DMRT3, DNAJA1, FANCC, FANCG, HEXB, KIF2A, LHFPL2, RAD23B, RPS6, SKP2, SPIN1, ZNF366*

*$ AR, DACH2, DIAPH2, MSN, SEPT6*

*& FSHR, LBH, MEA1, MSH2, LHCGR (NA in zebra finch)*

*€ AMD1, ARID4B, CCR6, CGA, CITED2, CYP1B1, ESR1, FOXO3, HSF2, LATS1, MAP3K4, MCM9, MEI4, NA (Uncharacterized protein), PACRG, PGM3, QKI, ROS1, RWDD1, SLC22A16, SRD5A2, STRN, TBPL1, TCF21, TCTE1, UBE2J1, UBR2, UFL1*

*! CELF1, EIF2B2, SLIRP, TTLL5, TYRO3*
